# *In vitro* and *in vivo* evidences propound therapeutic potential of Lipocalin 2 in cervical carcinoma

**DOI:** 10.1101/2023.01.13.523914

**Authors:** Nehanjali Dwivedi, Tahmina Mazumder, Gayathri Veeraraghavan, Ramanujam Siva, P K Smitha, Rohit Ranade, Manjula Das, Sujan K Dhar

## Abstract

Cervical cancer (CC), the second most common in developing countries and the third most common in developed nations, is the fourth most common type of cancer in women overall. The HPV16 high-risk genotype of the virus, which is responsible for about 61% of cervical cancer incidences, was found to have higher LCN2 levels in advanced clinical CC stages. In this study, we assessed the impact of suppressing LCN2 activity after treatment with an anti-LCN2 monoclonal antibody (MAb) in both *in vitro* and *in vivo* settings. Anti-LCN2 antibody was found to reduce proliferation and invasion of HeLa cells, the first immortal cells from a HPV positive aggressive adenocarcinoma of the cervix. LCN2 and its ligand MMP9 was found to be highly expressed in the cells and abrogated on treatment with anti-LCN2. The five receptors of LCN2 - SLC22A17, MC1R, MC2R, MC4R and LRP2 were barely detected with or without treatment. Anti-LCN2 Mab caused tumors to regress and soften *in vivo*, in a xenograft mouse model. Analysis of histology images of the treated and untreated tumor established the necrotic capability of the therapeutic molecule explaining the regression and softening of the tumor. Differential gene expression analysis between untreated and treated tumor proved that LCN2 inhibition abolished the migratory, invasive, and hypoxic pathways while significantly increasing the necrosis and cell death pathways in tumor after treatment with the monoclonal antibody. LCN2 inhibition was shown molecularly to lead to tumor regression via a negative feedback loop of LCN2 through the TNFα-IL17 axis exponentially increasing the effect of the anti-LCN2 monoclonal antibody. In conclusion, LCN2 appears to be a viable therapeutic target, and the monoclonal antibody used in this study can be further developed for clinical usage in cervical cancer.

## Introduction

The fourth most frequent form of female cancer overall, second in developing nations and third in developed nations, cervical cancer (CC) has a significant impact on women’s health all over the world^1^. India sees about 95000 – 1 lakh new cases every year^1^. According to the latest cancer statistics analysis in India, cervical cancer has the second highest incidence and mortality rate of 18 and 11.4%, respectively^1^. Unfortunately majority of cases are still diagnosed in advanced stages leading to death due to cervical cancer every 8 minutes in India.

It is mainly caused by high-risk human papillomavirus (HPV) infections^2,3^. The standard of treatment for early stage cancers (I-IIA) is a radical surgery (Radical hysterectomy with bilateral pelvic node dissection) followed by adjuvant radiation and chemotherapy according to the risk factors. While for locally advanced (IIB-IIIB) it is a combination chemoradiation. The treatment strategies/modalities depend upon the age, overall health of the patient and the cancer stage^4^. Overall for upfront metastatic and recurrent cervical cancers the treatment options are not very encouraging ^4^. A few monoclonal antibody treatments, such as tisotumab, bevacizumab and pembrolizumab have also been approved as a second line of therapy along with chemotherapeutic drugs. Although mainly caused by HPV infections, large number of cellular proteins are also known to be involved in promoting cervical carcinogenesis; for example various cytokines^5,6^, with a special mention about lipocalins, which have been recently reported to be involved in various human malignancies^7–9^. Even though majorly denoted as transport proteins, with respect to cancer, one particular member of the Lipocalin family, Lipocalin 2 (LCN2), has been particularly interesting^10^.

LCN2, also known as neutrophil gelatinase-associated lipocalin (NGAL) is a protein that in humans is encoded by the LCN2 gene ^11–14^. Human LCN2 was initially isolated and purified by Kjeldsen and coworkers as a 25-kDa neutrophil protein, that is in part associated with gelatinase from human neutrophils^15^. Very high levels are observed in the bone marrow^16^. LCN2 or NGAL is associated with innate immunity by sequestrating iron that in turn limits bacterial growth^17^.

Overexpression of LCN2 has been reported to increase the invasive potential of human cervical cancer ^18,19^. SiHa, a cervical cancer cell line reportedly increased its proliferative potential upon overexpression of LCN2 as demonstrated by Wang et.al ^19^. LCN2 levels were reported to be higher in advanced clinical CC stages with HPV16, a high-risk genotype of the virus that accounts for around 61% of cervical cancer incidences^20–23^, however its higher expression does not predict the incidence of cervical neoplasia^21^. Women with stage IV CC had a considerably higher quantity of LCN2 than women with stage I CC, indicating that LCN2 expression may be exploited to identify individuals with advanced disease who need more aggressive treatment ^23^. LCN2 levels were also found to be dependent on the duration of radiotherapy in patients, wherein, expression of LCN2 was significantly increased post ≥ 8 weeks of treatment with external beam radiotherapy (EBRT) and supplementary radiotherapy. However, in another study, it was demonstrated that lower LCN2 expression along with higher nm23-H1 expression correlated with poor prognosis and overall survival of the patients^24^.

In the current study, we evaluated the effect of abrogating LCN2 activity following treatment with an anti-LCN2 monoclonal antibody (MAb) in both *in vitro* and *in vivo* conditions. *In vitro* effects were assessed in HeLa, the first immortal human cell, “cancer in a test tube”, from a very aggressive glandular carcinoma of the cervix^25^. The molecular mechanism of action was predicted by qPCR on the cells and verified by end-point assays like migration, invasion and gelatine zymography. On the other hand, investigations in a tumor-induced transgenic animal model including tumor size regression, histopathological observation of tumor tissues, texture analysis of histopathology images and molecular profiling by transcriptome sequencing substantiated the *in vivo* efficacy. Our findings proclaim the tumor-regressive role of the anti-LCN2 monoclonal antibody and potentiates its therapeutic usage in cervical cancer to improve patient prognosis.

## Materials and Methods

### Analysis of public databases

The National Human Genome Research Institute (NHGRI)^26^ and the National Cancer Institute (NCI)^27^ have funded the Cancer Genome Atlas (TCGA)^28^, a project that has produced thorough, multidimensional maps of the major genetic alterations in many forms of cancer. Expression of the genes of interest were assessed in TCGA Cervical Cancer (CESC) dataset using RNA-seq expression values acquired from Broad Institute Firehose portal^29^ to compare their expression in tumor and normal samples.

### Recombinant LCN2 protein and anti-LCN2 monoclonal antibody production

Full-length, codon optimized human LCN2 cDNA (NM_005564) was synthesized and cloned by GeneScript, USA, between NdeI and XhoI restriction sites of pET-28b(+) vector, expressed in E.coli Rosetta cells either as a HIS-tagged protein or a tag-less protein. The HIS-tagged protein was purified on Ni-NTA (cat. no. 6562; BioVision; abcam, Inc.) column. The tag-less protein was purified using ion exchange chromatography. For hybridoma generation, BALB/c mice were injected with 10μg of purified LCN2 protein with Freund’s complete adjuvant in 1:1 ratio, followed by two boosters with a gap of 21 days each with Freund’s Incomplete Adjuvant in 1:1 ratio and a final injection with 50μg of protein prior to fusing splenocytes with Sp2/O cells. Monoclonal antibody was then isolated from the cell supernatant using Protein A beads from the single cell cloned antibody-producing colonies obtained through HAT/HT selection.

### HeLa cell culture

HeLa cells were cultured in DMEM medium, pH 7.2, (cat. no. 11995-065; Gibco; Thermo Fisher Scientific, Inc.) supplemented with 10% FBS and 1X penicillin-streptomycin solution at 37°C. The cells were further confirmed for their authenticity by STR profiling (Supplementary Table 1). STR multiplex assay was performed by TheraCUES, Bangalore using GenePrint 10 (Promega Corporation), version 3.0.0 of the SoftGenetics. GeneMarker_HID was used in order to analyze the result and the data was examined by referring to the STR Database of ATCC and CLASTR. For assessing the therapeutic potential of LCN2 as a target, tumorigenecity of HeLa cells was determined under the effect of anti LCN2 antibody as described in next sections.

### Proliferation Assay

HeLa cells were seeded at a concentration of 5×10^3^ cells per well in a 96-well plate, overnight at 37°C in 5% CO_2_ followed by a 12 h treatment with anti-LCN2 MAb and mIgG at indicated concentrations. Proliferative potential of the cells was calculated using Alamar Blue Reagent (cat. no. R7017; Sigma-Aldrich; Merck KGaA) at a final concentration of 0.2mg/mL, after 2 h of incubation at 37°C. Results show the mean standard deviation of relative fluorescence intensities of three independent experiments.

### Invasion Assay

Transwell inserts (cat. no. TCP083; HiMedia Laboratories, LLC) were coated with 100 μl of 1mg/mL ECM gel (cat. no. E1270; Sigma-Aldrich; Merck KGaA) in serum free (SF) medium followed by incubation for 2 h at 37 °C. HeLa cells (5×10^3^) were seeded in SF medium with indicated treatments of anti-LCN2 antibody and mIgG and allowed to invade for 24 hours. Following invasion, the inserts were stained with 2% crystal violet (cat. no. S012; HiMedia Laboratories, LLC) and quantified by eluting the stain with 10% acetic acid (cat. no. Q21057; Qualigen; Thermo Fisher Scientific, Inc.) as described before^30^. The findings are presented as the mean standard deviation of three independent experiments.

### Gelatin zymography

Gelatin zymography was performed by co-polymerizing the gels (SDS-PAGE, 10%) with gelatin (1 mg/ml) (cat. no. GRM019; HiMedia Laboratories, LLC). Ten microliters of gel-loading buffer and 20 microliters of concentrated conditioned medium was loaded onto the wells of the gel. Electrophoresis was carried out at a constant voltage of 125 V, until the dye reached the bottom of the gel, in cold conditions. After electrophoresis, gels were washed in washing buffer [2.5% Triton X-100 (cat. no. 10655; Fisher Scientific, Inc.) in 50 mM Tris-HCl (pH 7.5) (cat. no. T5941; Sigma, Inc.), 5mM CaCl_2_ (cat. no. C1016; Sigma, Inc.), 1μM ZnCl_2_ (cat. no. 28785; Fisher Scientific, Inc.)] for 1 h at room temperature with gentle agitation. The zymograms were then incubated for 18 h at 37°C in developing buffer [5 mM CaCl_2_, 1μM ZnCl_2_ in 50 mM Tris-HCl (pH 7.5), 1% TritonX-100]. Gels were then stained with Coomassie blue (cat. no. MB153; HiMedia Laboratories, LLC) and destained with 30% methanol (cat. no. AS059; HiMedia Laboratories, LLC) and 10% acetic acid. Gels were acquired and photographed by gel imaging system (Molecular Imager® Gel DocTM XR; Bio-Rad Laboratories).

### Animal husbandry

Male Ncr nude mice used in the study were housed in individually ventilated cages covered with stainless steel grid top. Temperature and humidity were maintained in the range of 19-23°C and 30-70%, respectively with 12-15 air changes/h. Animal confinement was provided with 12h artificial light photoperiod and 12h darkness. Animal experiments were performed under the approval of Institutional Animal Ethics Committee.

### Xenograft cervical tumor model

HeLa cells (5×10^6^) were injected using a needle of size 26 gauge subcutaneously (s.c.) into the dorsal flanks of NCr transgenic male nude mice. Mice were maintained under sterile and controlled conditions. Tumor growth was measured in 2 dimensions with a caliper using the formula (0.5x LV^2^) where L is the largest dimension and V its perpendicular dimension. Once tumors grew to about 100mm^3^ in diameter, tumors were injected with anti-LCN2 antibody every alternate day (100μg) directly into the tumor with PBS as the vehicle control in no treatment group. Animals were sacrificed when the tumor grew °2cm^3^, after 2 weeks of treatment.

### Histolopathology analysis

Animals were sacrificed and tumors were removed after 2 weeks of treatment with anti-LCN2 antibody. Half of the tumors were fixed in 10% (v/v) formalin (Cat. no. HT501128; Sigma) and embedded in paraffin (cat. no. Q19215; Qualigen; Thermo Fisher, Inc) for sectioning (n=3 in each group). Sections of 5μm thickness were stained with hematoxylin and eosin. Slides were analyzed for the extent of tumor boundary and regions of necrosis and other cellular events before converting to digitized images using a Grundium ocus40 slide scanner.

### Texture analysis of histopathology images

H&E-stained histopathology images of treated and untreated tumor samples were analyzed to extract some of the grey-level co-occurrence matrix (GLCM) features originally proposed by Haralick et al^31^. Whole-slide images were tiled into 16 × 16 pixel patches with minimum 90% tissue content and colour normalized. Three basic GLCM features including entropy, contrast and dissimilarity which are measures of randomness and disorder in the images were calculated for each image patch using python scikit image library. For a visual representation, the feature value of each patch was converted to a colour map with red to green colour gradient indicating maximum to minimum value of each feature.

### RNA isolation, cDNA conversion and qPCR

RNA isolation from 1×10^6^ cells was performed using TRIzol method, as per the manufacturer’s instructions. Briefly, 1×10^6^ cells were resuspended and incubated for 10 minutes on ice in 1mL of TRIzol^32^ followed by addition of 200μL of chloroform and centrifugation at 13,000 g for 15 minutes at 4°C. The aqueous layer was transferred into a fresh tube containing equal volume of 100% isopropanol without disturbing either the middle interface or the lower organic phase. The tube was gently invert-mixed and incubated at room temperature for 10 minutes followed by centrifugation at 13,000 g for 10 minutes at 4°C. The supernatant was discarded and the pellet was washed with 75% ethanol followed by centrifugation at 7500 g for 5 minutes at 4 °C. Air-dried pellet was then dissolved in nuclease free water for further processing and 1μg of total RNA was used for cDNA conversion using AMV Reverse Transcriptase (NEB, USA) in a 20μl reaction. For qPCR a set of two reference genes - ACTB and GAPDH ^33,34^ were used in the study. On a Roche LightCycler 480 II instrument using KAPA SyBr green Universal, qPCR was performed in triplicates. In a total of 5μl reaction volume containing SyBr mix, 1:5 diluted cDNA template, primers (100nM each), and water were used. The reaction conditions were: Pre-incubation at 95°C for 10 seconds followed by 45 cycles of amplification (95°C – 1 second; 95°C – 10 seconds; 60°C – 15 seconds; and 72°C – 15 seconds). List of primers are detailed in supplementary table 2.

### Transcriptome analysis

RNA was isolated from ∼ 1cm^3^ of the tumor using Qiagen RNA isolation kits (Cat. No. 74004; Qiagen) as per manufacturer’s instructions. Library for RNAseq was prepared using illumina TruSeq library preparation kit and sequencing was performed on a NovaSeq 6000 (illumina) platform using 2 × 150 bp chemistry at LifeCell International, India. Sequence reads were filtered for quality using Trimmomatic ^35^ and filtered reads were separated based on mapping to human (GRCh38) and mouse (GRCm39) reference assembly using FastQ Screen^36^. Separated reads were aligned to respective genome using STAR pipeline ^37^ and differentially expressed genes were determined using edgeR^38^. Functional and pathway analysis of the differentially expressed genes were carried out using EnrichR web server ^39^ and ClusterProfiler R library ^40^. Gene expression heatmaps were drawn using pathway signatures from MDSigDB ^41^ and published literature.

## Results

### Expression of LCN2 and ligands in Cervical Cancer

LCN2 is known to have intracellular effect through its receptors in a cell specific manner^42,43^ or an extracellular effect by binding to MMP9 and stabilizing it, thereby increasing the protease activity^44^. Analysis of expression of LCN2 and its known ligands MMP9 and the five receptors in the TCGA public database revealed that LCN2 and MMP9 have a significantly higher expression in cervical cancer tumors as compared to normal samples (figure 1a,b), whereas SLC22A17, the most widely known LCN2 receptor showed the opposite behaviour (figure 1c). All the other receptors did not show any significant difference between normal and tumor samples, while MC1R remained undetected. In house analysis using qPCR in HeLa revealed the same pattern, wherein, LCN2 and MMP9 are overexpressed as compared to the 5 receptors (figure 1b), thereby confirming only MMP9 mediated action of LCN2 on HeLa cells.

**Figure 1.**
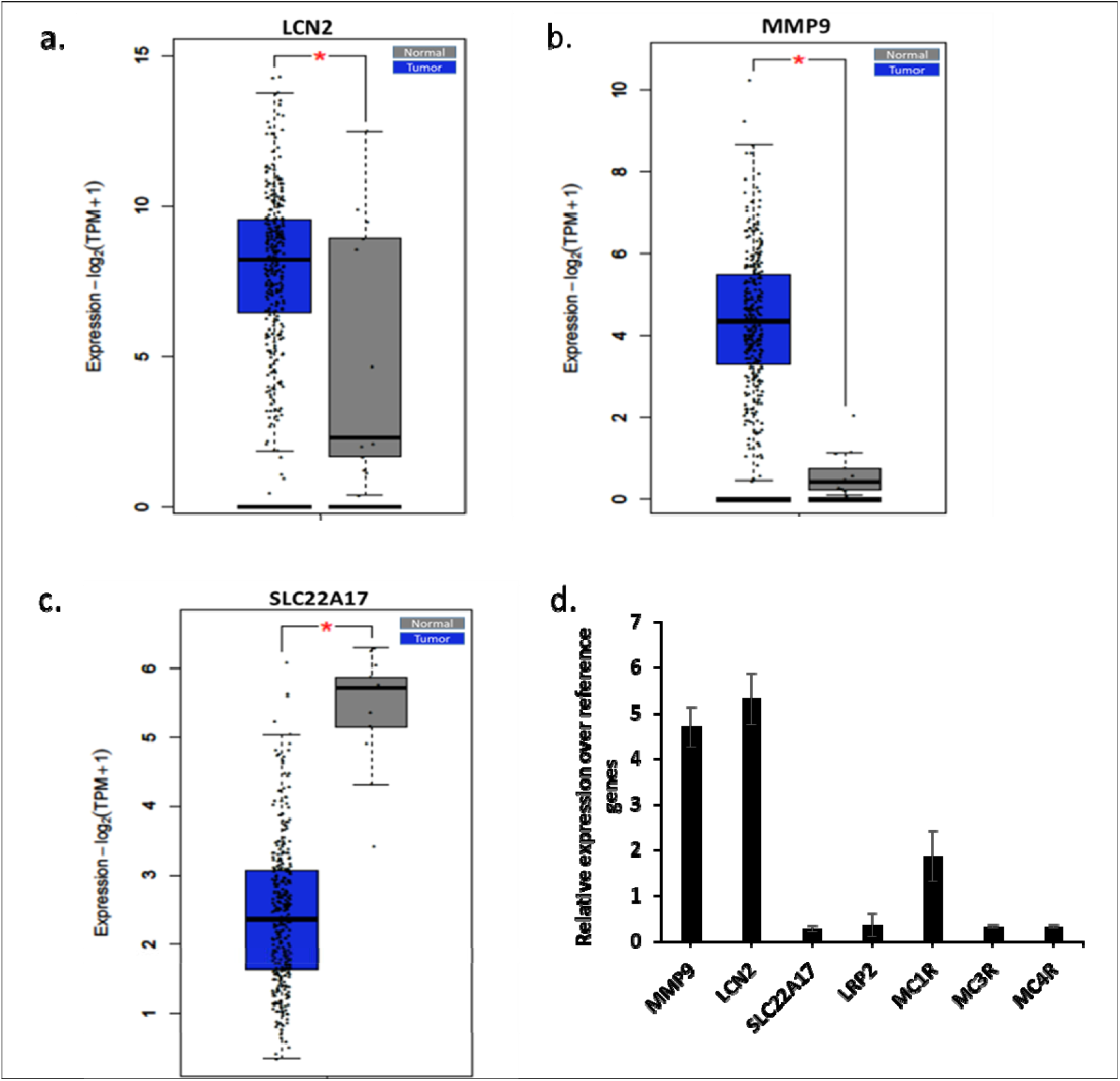
mRNA expression analysis. (a) LCN2 expression in TCGA database between tumor and normal samples, (b) MMP9 expression in TCGA database between tumor and normal samples, (c) SLC22A17 expression in TCGA database between tumor and normal samples, (d) qPCR analysis of expression of indicated genes on HeLa cells

### Anti-LCN2 antibody abrogates the tumorigenicity of cervical cancer cell line

Treatment with anti-LCN2 antibody decreased the proliferative ability of HeLa in a dose dependent manner starting as early as 12 hours (figure 2a). Similarly, a significant increase in the inhibition of proliferation in the same manner as depicted by dashed line over the proliferation graph (figure 2a). Treatment with anti-LCN2 antibody reduced the invasive ability of the cells in a dose dependent manner (figure 2b,c). Control mIgG addition did not show any statistical significance at the equivalent highest molar concentration of anti-LCN2 antibody, suggesting that the decrease in tumorigenicity is specifically due to anti-LCN2 antibody.

**Figure 2.**
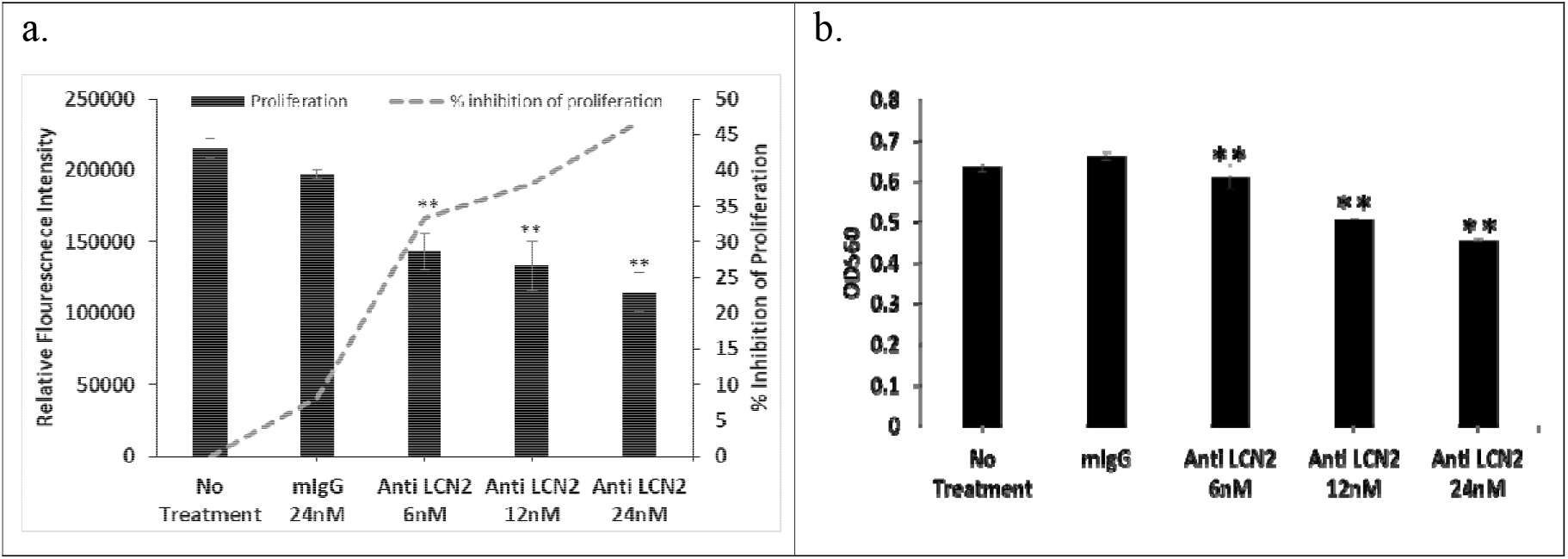
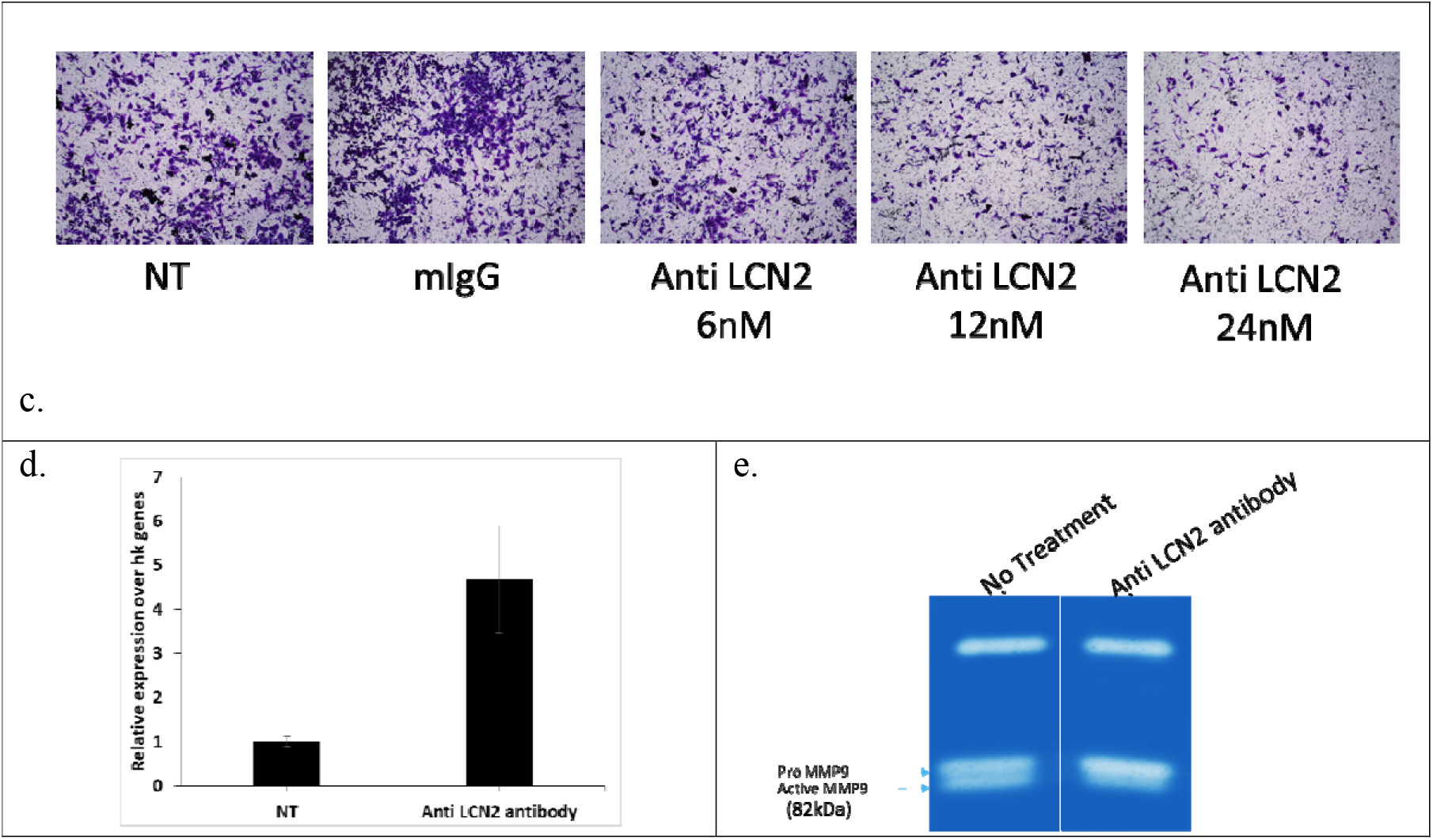
Effect of treatment of anti-LCN2 antibody on HeLa cells. (a) Proliferative potential as measured by alamar blue, (b) Quantitative representation of the invaded cells under the effect of anti-LCN2 antibody, (c) pictorial representation of invaded cells under the effect of anti-LCN2 antibody stained with crystal violet. (d) qPCR expression of Claudin 1 in HeLa cells upon treatment with anti-LCN2 antibody (e) Gelatin zymography to check the MMP9 activity upon indicated treatments. All images represent 4X magnification. **p<0.05.

Claudin is a tight junction protein, which is known to get downregulated when cells undergo migration^45^. Expression of Claudin mRNA increased upon treatment with anti-LCN2 antibody suggesting the restriction in the migratory abilities of the cells under the action of anti-LCN2 antibody (figure 2d). MMPs, well-known metalloproteases are involved in invasion and matrix degradation. MMP9, one of the metalloprotease has also been shown to form complex with LCN2, which further stabilizes MMP9 and its activity^46^. HeLa cells upon treatment with anti-LCN2 antibody, showed a reduction in the expression of active metalloproteases in gelatine zymography experiment, suggesting a quick degradation of these proteases and thus inhibiting migratory and invasive ability of HeLa cells (figure 2e). Molecular weight of the clearing zone indicates the protease to be MMP9.

All the data taken together suggests that LCN2 abrogation resulted in inhibition of migratory and invasive potential of HeLa cells.

### Tumor Formation

Tumor formation was observed at the site of injection after approximately 7 days of subcutaneous HeLa cell line administration. During the time of treatment, animals appeared to be normal without any distress but for the tumor related distress. After reaching a volume of 100mm3, the tumors were treated with anti-LCN2 antibody at a concentration of 100μg/mL every alternate day, directly into the tumor along with the vehicle control in the no treatment group. After 2 weeks of treatment, once the control group’s tumor grew to ∼200 mm^3^ in size, all the animals were sacrificed because of ethical considerations.

### Effect of Anti-LCN2 on xenograft

As seen from figure 3a, regression in tumor could be observed as soon as day 6 of the treatment with anti-LCN2 antibody. Histopathological analysis revealed that the control group showed tumor epithelial cells with clearly defined boundaries as shown by purple arrow in figure 3b. Tumor sections obtained post treatment revealed areas of necrosis and fibrosis demarcated by the tissue areas having shrivelled cellular morphology. Such areas are marked in yellow arrows in figure 3b. Treatment with anti-LCN2 antibody thus showed tumor regression characterized by necrotic regions in the tumor, promulgating the therapeutic potential of LCN2.

**Figure 3.**
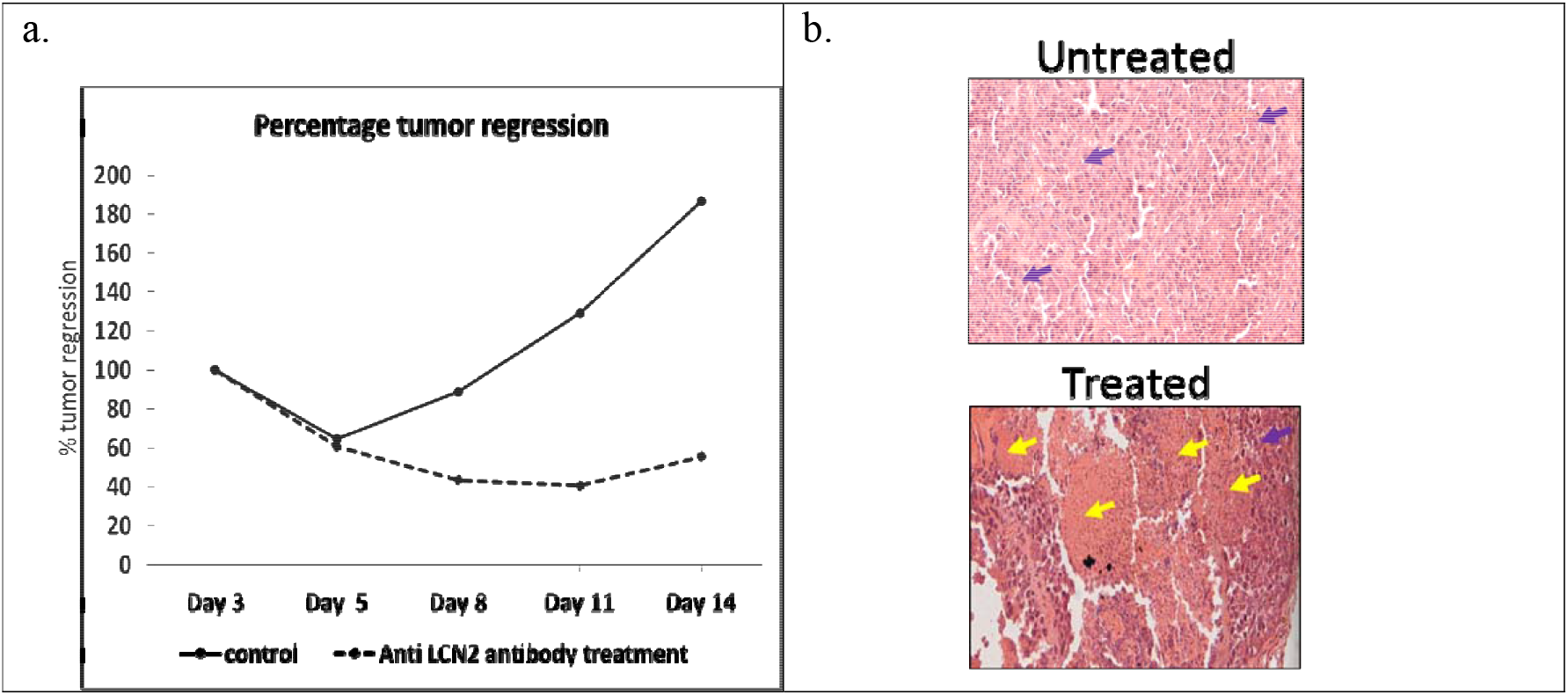
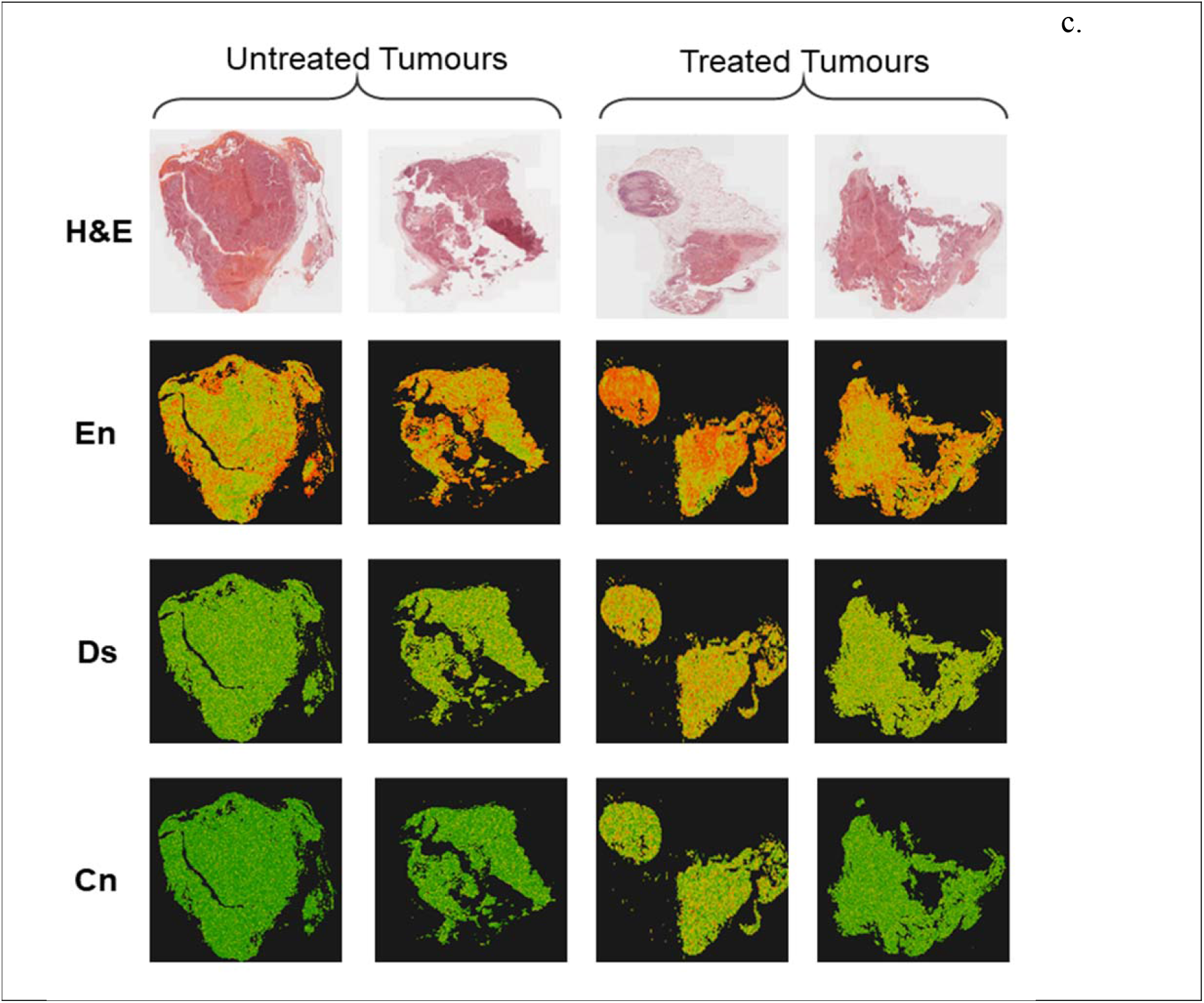
Effect of anti-LCN2 antibody on tumor regression. (a) Percentage tumor regression post treatment with anti-LCN2 antibody, (b) Histological analysis of the tumor obtained from control group, and the treatment group.(c) H&E-stained histopathology images of intreated and treated tumors and corresponding visual representation of entropy (En), dissimilarity (Ds) and contrast (Cn) values for each 16×16 px in the images using a a colourmap with red to green colour gradient indicating maximum to minimum value of each feature. More red dots are seen in treated sample images signifying higher degree of disorder.

Texture features of whole-slide histopathology images such as entropy, dissimilarity and contrast have been correlated in literature with clinical end-points or molecular subtypes in multiple cancers including breast^47^, oral^48^ and multiple other types of cancers^49^. In this analysis, the three features - entropy, dissimilarity and contrast, presented with higher values in images from treated relative to untreated tumors (figure 3c) indicating emergence of disorder due to necrosis of cells in treated samples. The difference in each parameter was found to be statistically significant (p << 0.001) using t-test. The same was independently confirmed by histopathologist as indicated in figure 3b.

### Tumor regression and molecular mechanisms

Since the tumor was formed in another host organism, i.e. nude mice, sequencing reads were segregated into mouse and human components based on mapping to reference assemblies (figure S1a). Unique human fragments mapped were more in number as compared to the unique mouse fragments evidencing human origin of extracted tumor. Mouse genes differentially expressed in treated samples revealed activation of cytokine pathways due to introduction of foreign human cells in the host (figure S1b).

Enrichment analysis of differentially expressed genes revealed that some of the major players in IL17 signalling pathway including NFkB and CXCLs are downregulated on treatment (Figure 4a). Amongst the cancer hallmark genesets, TNF-α signalling pathway genes were found to be downregulated upon treatment (figures 4b and 4c). LCN2 has been reported to be a downstream effector molecule of IL17 signalling pathway, enhancing the tumorigenicity of gastric cancer^50^. Moreover, overexpression of LCN2 has been associated with diagnosis of late stage cervical cancers^23^. LCN2 has also been reported to enhance tumor cell migration and invasion both in vitro and in vivo in cervical cancers via stimulation of EMT pathway^18^ In addition, LCN2 was also observed to be downregulated in the treatment group, suggesting a negative feedback loop mechanism of antitumor efficacy of anti-LCN2 antibody, i.e. anti-LCN2 antibody abrogates the activity of LCN2 in tumors, which in turn downregulates the IL17 signalling pathway, further downregulating LCN2 in the process, thus exerting its anti-tumor activity.

**Figure 4.**
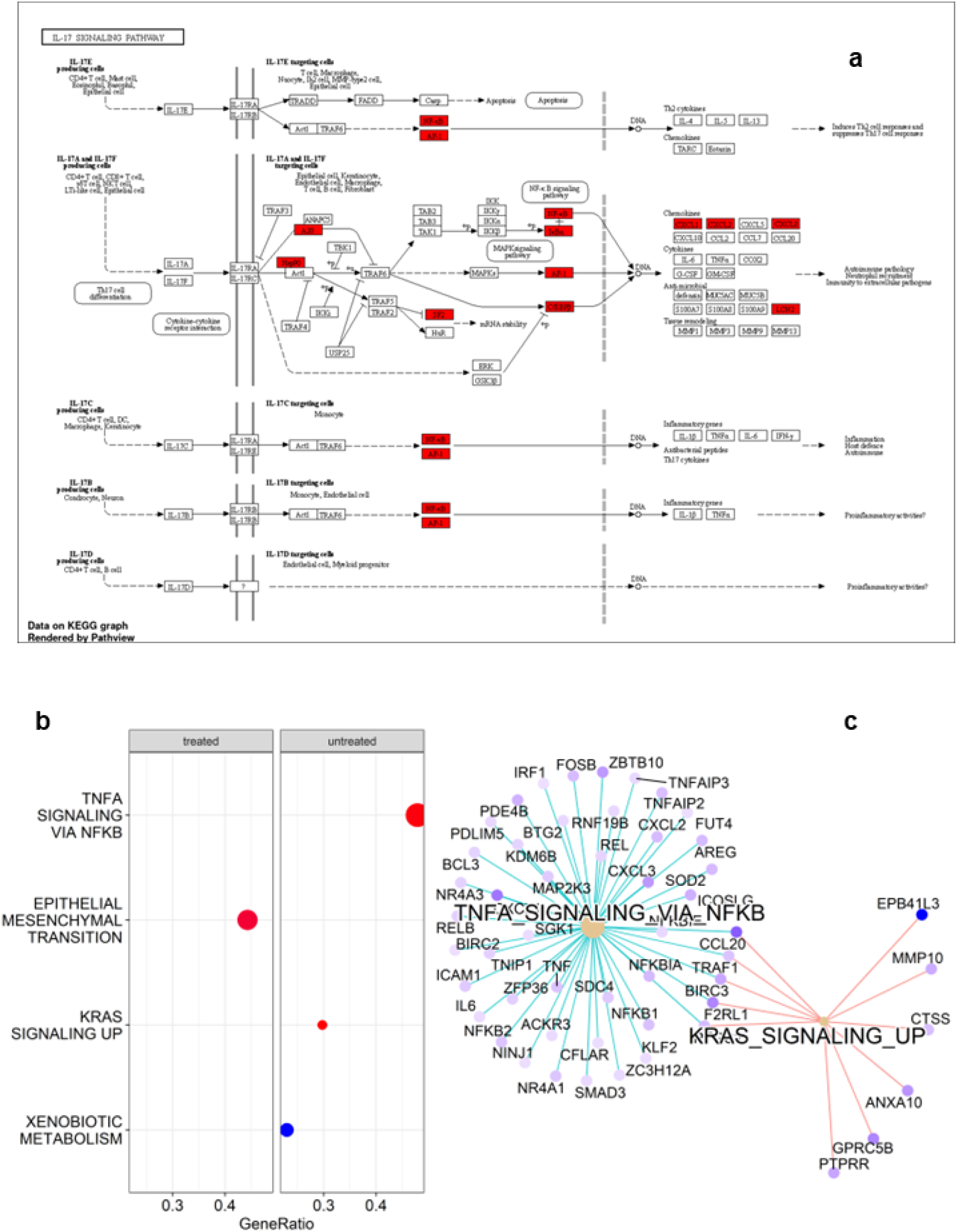
Functional analysis of differentially expressed genes depcting (a) Genes (shaded in red) downregulated in the treatment group, (b) cancer hallmark genesets enriched in treated and untreated samples and (c) genese associated with TNF-α and KRAS signalling pathways enriched only in untreated samples (and hence suppressed under treatment).

This was in concordance with our expectation as the tumor was treated with anti-LCN2 antibody. Further, pathway analysis of the differentially expressed genes revealed that the treated samples had higher expression of genes pertinent to cell death pathways (cell death pathway gene signature taken from GSEA mSigDB^41^) (figure 5a), which was again in concordance with the histopathological analysis performed. These results thus suggest that anti-LCN2 antibody treatment increased the cellular death pathways in the treated tumor, thus promoting fibrosis and necrosis of the same, a pre-requisite of tumor regression. Upon further analysis of the tumorigenicity pathways from literature, it was observed that the untreated samples had higher expression of genes involved in promoting invasion (figure 5b), hypoxia (figure 5c), and migration (figure 5d).

**Figure 5.**
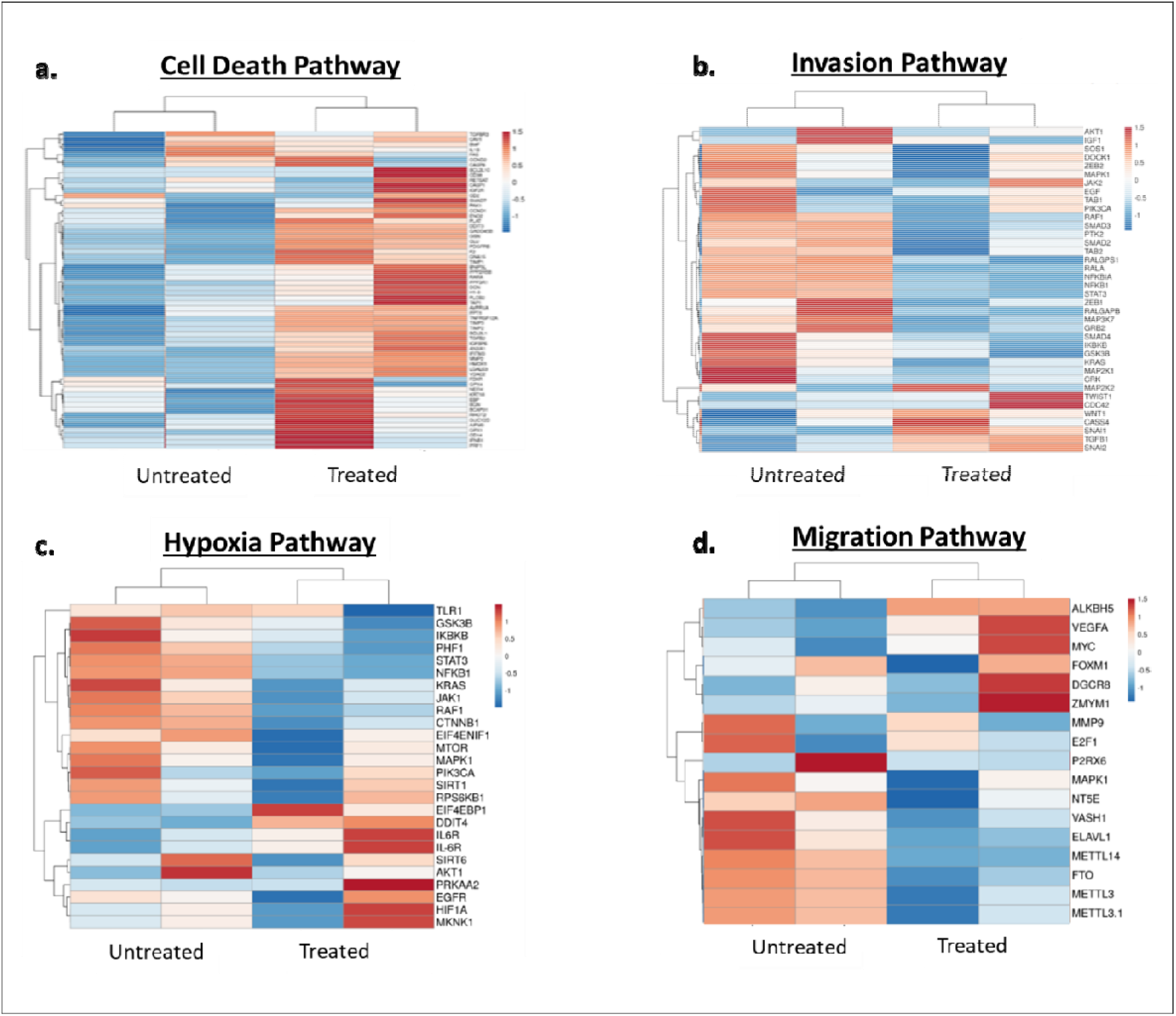
Heatmaps showing pathway expression between untreated and treated samples. (a) Cell death pathway, (b) Invasion pathway, (c) Hypoxia pathway, (d) Migration pathway

## Discussion

LCN2 is known to be a potential therapeutic target in a large number of solid human malignancies. A mesenchymal-like cell morphology has been reported to be induced by LCN2 overexpression, along with overexpression of various mesenchymal markers (Snail, Twist, N-cadherin, fibronectin, and MMP9) that promote invasiveness, thus explaining the role of LCN2 overexpression in tumor cell motility during metastasis ^51^.

In a report by Wang and colleagues ^24^, it was demonstrated that nm23-H1 knockdown mediated increase in LCN2 promoter activity resulted in decreased cell migration and invasion in SiHa uterine cervical cancer cells. Comparing cervical carcinoma to high- and low-grade dysplasia tissue, LCN2 expression is actually higher in the latter ^52^. Additionally, Snail, Twist, N-cadherin, fibronectin protein levels and MMP-9 activity increased in C33A cells when LCN2 was overexpressed. Core transcription factors in the EMT process, the mesenchymal markers Snail and Twist have been reported to activate the downstream target genes encoding N-cadherin, fibronectin, and MMP-9 under the effect of LCN2 ^53,54^.

In this study, inhibiting LCN2 using anti-LCN2 antibody resulted in suppression of major tumor promoting signalling pathways like TNF alpha, IL17 and NFkB signalling, which ultimately resulted in tumor regression. Numerous reports suggest that all these pathways are linked to LCN2 in various human malignancies^55–59^. For example, in PC-3 prostate cancer cells, it has been reported that TNF alpha stimulates LCN2, thus protecting the tumor cells from apoptotic cell death. Reports have also suggested that TNF alpha is directly associated with susceptibility to cervical cancers ^60^. IL17 has also been implicated to have a tumor promoting role in cervical cancers via activation of NFkB pathway ^44,61,62^. Downregulation of these tumor promoting pathways by treatment of anti-LCN2 antibody in the xenograft model in this study proves conclusively that LCN2 can be a potential therapeutic target for abrogating tumor progression.

Furthermore, during the study it was observed that the tumors of the treatment group became softer as compared to the control group, which still remained very stiff and packed. Increased tumor stiffness is a widely approved characteristic of solid tumors^63^. Piao et.al reported in cervical cancer as well that the substrate stiffness affected the epithelial to mesenchymal transition of HeLa and SiHa cervical cancer cell lines^64^.

Identification of cytokine dysregulation has been of crucial significance in the development and spread of cervical cancer ^65,66^. Specifically IL17 has been regarded as a significant factor in the progression of various human malignancies, including cervical cancer. Bai et.al showed that IL17 regulates the progression of cervical cancer via the activation of JAK/STAT3 and PI3K/NFkB pathway^61^. Other reports also suggest its involvement in the upregulation and activation of MMP2 as well as MMP9, making it a potential target to improve prognosis for patients diagnosed with cervical cancer^44,62^. Further to this, Tartour et.al suggested that IL17 overexpressing cervical cancer cell lines promoted tumor formation in nude mice^67^. Additionally, IL17 has been implicated in the progression of cervical carcinogenesis, not only with high-risk HPV infection^68^, but also with HPV negative cervical cancer cells^68^.

Recent reports have also suggested that matrix stiffness plays an important role in the prediction of aggressiveness and affects the morphology, drug sensitivity and mechanical properties of cervical cancer cells^69–71^. It can therefore be stated that treatment with anti-LCN2 antibody decreased the stiffness of the tumor, thus helping in tumor regression. The softening of tumor by anti-LCN2 administration opens an avenue for localized treatment by intra-tumoral injection. However, more mechanistic experiments have to be performed to correlate the involvement of LCN2 in decreasing tumor stiffness.

In conclusion, it can be stated that LCN2 inhibition improves tumor regression properties both in vitro and in vivo. By encouraging cancer cell motility by activating EMT pathway components, LCN2 appears to be a potential therapeutic target. The monoclonal antibody that abrogates the stiffness and invasive-migratory properties of the tumor can be developed further for clinical usage.

