## Supplementary Figures for "*In vitro* and *in vivo* evidences propound therapeutic potential of Lipocalin 2 in cervical carcinoma"

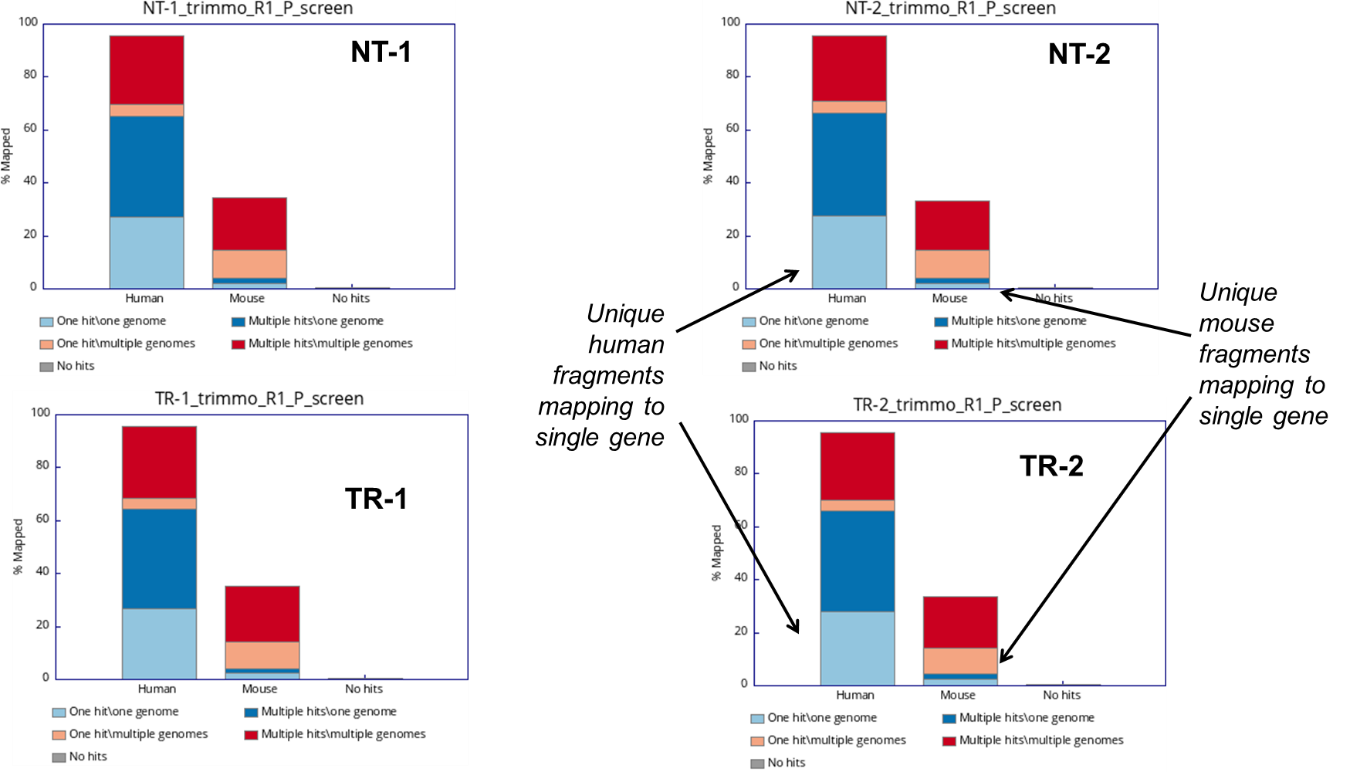
a.

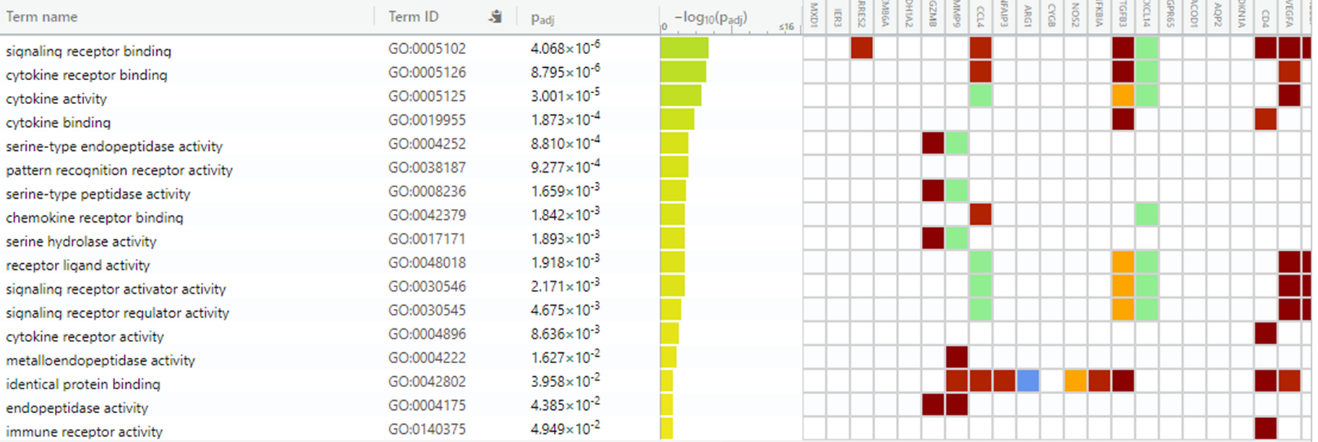
b.

Supplementary figure 1: Sequence analysis of the tumors. (a) Species identity of the sequencing sample, (b) Pathway association of the differentially expressed genes mapped to mouse genome.

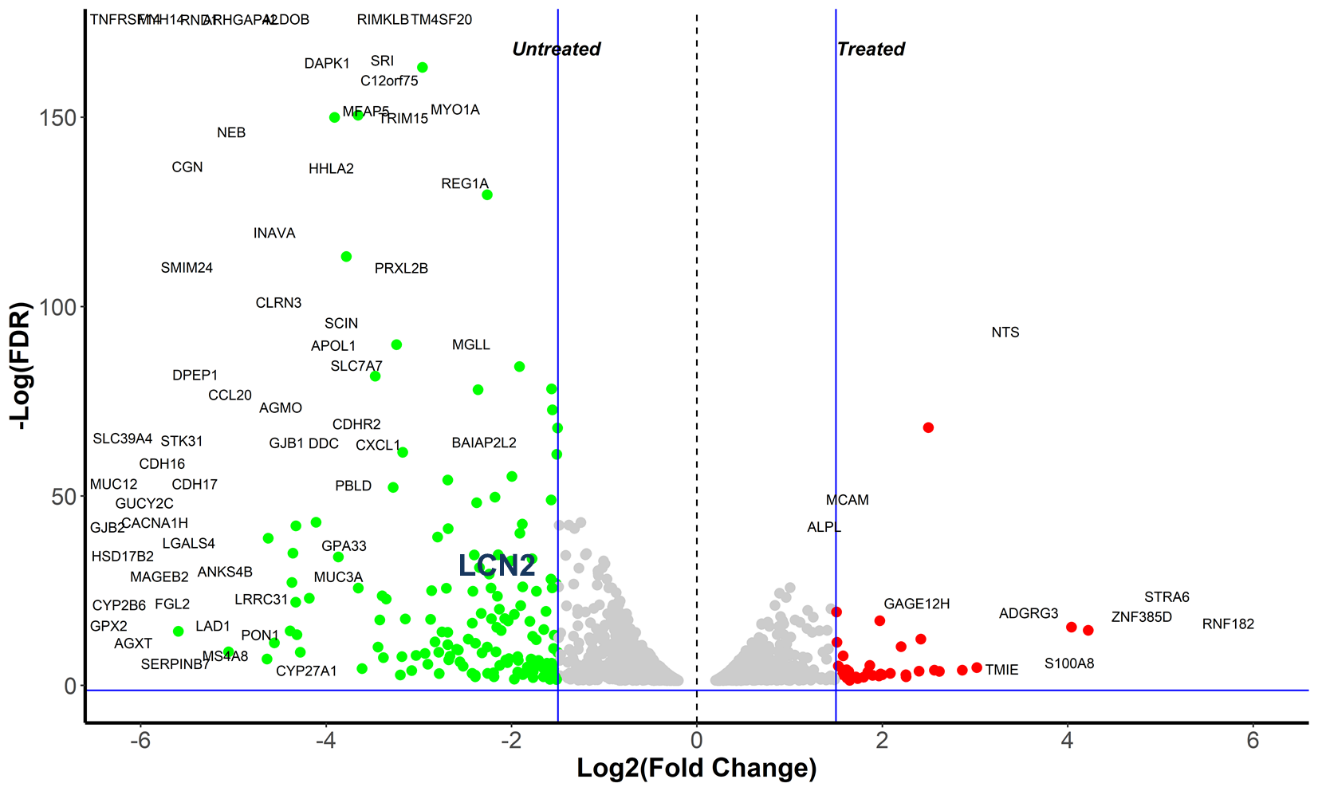

Supplementary figure 2 Differential expression analysis showing LCN2 expression in untreated samples.
