## Supplementary Tables for "*In vitro* and *in vivo* evidences propound therapeutic potential of Lipocalin 2 in cervical carcinoma"

Supplementary Table 1: STR analysis of HeLa cell line. The STR profile matched 100% to HeLa cell line from ATCC database ^1^ as authenticated by TheraCUES.

|  | Marker | Allele#1 | Allele#2 |
| --- | --- | --- | --- |
| **HLA** | TH01 | 7 | 7 |
|  | D21S11 | 27 | 28 |
|  | D5S818 | 11 | 12 |
|  | D13S317 | - | - |
|  | D7S820 | 8 | 12 |
|  | D16S539 | 9 | 10 |
|  | CSF1PO | 9 | 10 |
|  | AMEL | X | X |
|  | vWA | 16 | 18 |
|  | TPOX | 8 | 12 |

Supplementary Table 2 Primer sequences used in the study

| GAPDH | Fw 5'-TCGACAGTCAGCCGCATCTTCTTT-3'  Rv 5'-GCCCAATACGACCAAATCCGTTGA-3' |
| --- | --- |
| ACTB | Fw 5'- GCGCGATATTTCTTCTTGCAGG-3'  Rv 5'- TTCGTACCTGGCATGACTGG-3' |
| Claudin | Fw 5'-AGCCATGTACGTTGCTATCCA-3'  Rv 5'-ACCGGAGTCCATCACGATG-3' |
| SLC22A17 | Fw 5'- GTTACCCCGCAGACAGATTTG-3'  Rv 5'- CCCAAGAGGAATCGGAGGG-3' |
| LRP2 | Fw 5'- GTTCAGATGACGCGGATGAAA-3'  Rv 5'- TCACAGTCTTGATCTTGGTCACA-3' |
| MC1R | Fw 5'- CATCGCCAAGAACCGGAAC-3'  Rv 5'- GGATGACGGCCGTCTCC-3' |
| MC3R | Fw 5'- GGGCATCGTCAGTCTGCTG-3'  Rv 5'- AGGGCATTGGACACACTTACC-3' |
| MC4R | Fw 5'- ACACCCACTCCTCCACCTTT-3'  Rv 5'- TGCTGTAGCCAAATTCGTTG-3' |
| LCN2 | Fw 5'- GAAGTGTGACTACTGGATCAGGA-3'  Rv 5'- ACCACTCGGACGAGGTAACT-3' |
| MMP9 | Fw 5’ – TTGGTCCACCTGGTTCAACT – 3’  Rv 5’ – ACGACGTCTTCCAGTACCGA – 3’ |

References

1.	HeLa - CRM-CCL-2 | ATCC. https://www.atcc.org/products/crm-ccl-2.
